# Phenology and Robustness in plant-pollinator networks

**DOI:** 10.1101/2020.10.30.362616

**Authors:** Laura Melissa Guzman, Scott A. Chamberlain, Elizabeth Elle

## Abstract

Many metrics that describe the structure of mutualistic plant-pollinator networks have been found to be important for network stability and robustness. These metrics are impacted by a suite of variables, including species traits, species abundances, their spatial configuration, and their phylogenetic history. Here, we consider a specific trait, phenology, or the timing of life history events. We expect that timing and duration of activity of pollinators, or of flowering in plants, could greatly affect the structure of the networks in which they are embedded. Using plant-pollinator networks from 33 sites in southern British Columbia, Canada, we asked a) how phenological species traits, specifically timing of first appearance in the network and duration of activity in a network, were related to network structure, and b) how those traits affected network robustness to phenologically biased species loss. We found that long duration of activity increased connection within modules for both pollinators and plants and among modules for plants. We also found that date of first appearance was positively related to interaction strength asymmetry in plants but negatively related in pollinators. Networks were generally more robust to the loss of pollinators than plants, but robustness declined with loss of early-flying or long-duration pollinators. These pollinators tended to be among-module connectors. Our results show that changes in phenology have the potential to impact plant-pollinator networks, which may have conservation relevance in a time of changing climate.

## Introduction

Species within communities form interaction networks, and many metrics that describe the structure of mutualistic plant-pollinator networks (e.g. interaction asymmetry) have been found to be important for the ability of networks to be resilient to perturbations (Bascompte and Jordano 2007). Many species attributes contribute to network structure, including traits like flower colour and feeding preferences (Vázquez et al. 2009, Valdovinos 2019), abundance (Vázquez et al. 2007, Valdovinos 2019), spatial configuration (Morales and Vázquez 2008), and phylogenetic history (Rezende et al. 2007, Chamberlain et al. 2014). Other traits, such as phenology (the timing of life history events), have been less studied in the context of community resilience, even though timing can be crucial for finding a mate, provisioning a nest, and other fitness correlates.

Phenology is also important to consider in the context of perhaps the biggest perturbation communities will experience during our lifetimes: climate change. We know that phenology shifts with climate (Bartomeus et al. 2011), and there is some evidence that plants and pollinators are not responding similarly. For example, insect phenology is advancing more than plant phenology, and early-season angiosperms advance more than those that flower later in the season (Hegland et al. 2009, Wolkovich et al. 2012). Across 10 bee species in northeastern North America, phenology has advanced by ca. 10 days over 130 years (Bartomeus et al. 2011). This study also found that the plants from the same location and that flower during the same time as the pollinators are active have advanced at the same rate. Communities with higher biodiversity may be buffered against shifting phenologies because of varying responses and complementarity in activity periods of different species (Bartomeus et al. 2013b). However, given the rapid pace of climate change and its global scale, we need to understand how the phenology of specific species determines network structure, and the potential impact of shifting species phenologies on network structure and robustness. Specifically, we explored two variables the can describe the phenology of a species: when a species emerges or flowers for the first time during a season, and how long a species is active during the season. Shifts in the timing of both of these variables can lead to mismatches with potential interaction partners (Hegland et al. 2009), potentially affecting network robustness.

Network structure is associated with species traits. For example, modules in a network — groups of species that interact more with one another— can be the result of habitat heterogeneity, co-evolution or phylogenetic relatedness of the species (Pimm and Lawton 1980, Lewinsohn et al. 2006, Thompson 2005). In seed dispersal networks for example, plant and animal trait values —body mass and seed mass— were associated with the modularity of individual species (Donatti et al. 2011). Phenological traits have also been shown to be associated with modularity (Morente-Lopéz et al. 2018). Other network properties associated with species phenologies include interaction turnvover and rewiring (CaraDonna et al. 2017). In addition, species phonologies can be used to re-construct networks and the metrics describing those networks (Olito and Fox 2014).

Metrics that describe networks, such as modularity, have also been associated with the stability of networks (Thebault and Fontain 2010). Other aspects of network structure that are important for robustness are the degree of specialization of species and the asymmetry in the interactions (Kaiser-Bunbury et al. 2010, Mello et al. 2011). Given the relationship between species phenological traits and network properties, and the relationship between network properties and network stability, we would expect that phenological traits themselves would be the mechanism that underlies the relationship between network structure and network stability. Indeed, Encinas-Viso et al. (2012) and Ramos-Jiliberto et al. (2018) studied the effect of phenological traits on the stability and robustness of networks. Both of these studies used dynamical models to study this relationship. Encinas-Viso et al. (2012) found that as the length of the season in a network increases, diversity and resilience of the network also increases. Ramos-Jiliberto et al. (2018) went a step further and combined these dynamical models with empirical networks, and found that the loss of plants with earlier blooming dates and with longer active periods decreased pollinator persistence.

Here we use 33 mutualistic plant-pollinator interaction networks from Western Canada to ask how plant and pollinator phenology contribute to their network interaction structure. We focus on exploring four measures of network structure that are related to robustness: specialization, within-module degree, among-module connectivity, and interaction asymmetry. Specifically, we ask the following two questions: 1) How do date of first appearance in a network, and length of activity during the season, affect individual species interaction patterns? We predict that species whose date of first appearance is earlier, and that are active longer in the season, should be less specialized, have greater within-module degree, greater among-module connectivity, and have higher values of interaction asymmetry (they affect their partners more than the reverse). 2) How robust are networks to removal of species due to varying phenological “traits” (date of first appearance early/late, duration of activity small/large)? Networks should be more robust to losing species whose date of first appearance is later in the season, and species that are active during less of the season.

## Methods

### Study sites

A total of 33 mutualistic plant-pollinator networks were studied in British Columbia, Canada: oak savannah (12 networks), shrub-steppe (eight networks), and hedgerow restorations (13 networks). These three vegetation types comprised three different studies. See Table A1 for site information, including latitude/longitude coordinates. For simplicity we use “pollinator” throughout this paper to refer to insects and hummingbirds observed visiting flowers and contacting reproductive organs, although their effectiveness at transfer of pollen has not been assessed.

### Collection of mutualistic network data

Data were collected for two of three vegetation types using the plot method, and for the third using the transect method. Plots are generally more appropriate when the plant species in the community are very patchily distributed (Gibson et al. 2011), as they were in these regions. The plot method focuses on individual plant species, observing each plant species for an equal amount of time. For oak savannah sites (within the Coastal Douglas Fir biogeoclimatic zone), we collected data on species interactions in 1-ha plots at each of six sites in both 2009 and 2010. Each plot was surveyed about every 7-10 days, 10-12 times per season between late April and early July, the majority of the flowering period. Over the flowering period we attempted to visit sites morning, midday, and afternoon on different survey dates to reduce bias due to flight time differences among visiting insects. During each survey date, each plant species in flower was observed for a 10 min period by each of two surveyors, on haphazard walks throughout the plot. All flower visitors were collected and identified to the lowest taxonomic level possible. For the eight shrub-steppe sites (in the Bunchgrass biogeoclimatic zone), data were collected as for oak-savannah sites, but surveys were from the beginning of April through the end of July, 2010, for a total of 12 samples per site.

For hedgerow restoration sites, data were collected using a “transect” method, in which the plants along the transect were observed for a set amount of time, with time observed per plant species varying among species depending on their occurrence in the transect. Transects are more appropriate when plants are not clumped, but are widely scattered throughout a study site (Gibson et al. 2011), and in this case most of the restorations (within the coastal Western Hemlock biogeoclimatic zone) were linear, making transects efficient. Sampling was equal across all sites, occurring approximately every 2 weeks, for a total of 9 samples between late April and the end of August, 2013. The transect was walked for 15 minutes by each of two observers during each sample date, and each site was again observed equally during morning, midday, and afternoon on different sample dates. All pollinators were collected for identification in the lab.

### Species phenological variables

We collected the following phenological species variables for every pollinator and plant species: a) first Julian day observed interacting in the network, and b) number of days observed in the network (last date observed – first date observed). We treat each network as a replicate, and capture both variables for all species within the network. This means that there is no single value for first day observed or total days observed for any particular species across networks. Phenological variables were calculated from the observation of the interactions. While we acknowledge this is an imperfect way to detect phenological variables, it is consistent across all 33 networks and has been previously used in the literature (Rasmussen et al. 2013). Because these two phenological variables could potentially be correlated (they are calculated from the same set of data), we calculated Pearson’s correlation coefficient for log10-transformed variables for each of plants and pollinators separately. We found that the two variables were weakly negatively correlated (pollinators: ρ = −0.5, *P* < 0.001, df = 1690; plants: ρ = −0.25, *P* < 0.001, df = 590), meaning that if a species has a very late First Julian day it cannot, by definition, be present many days in the network. In contrast if a species is has an early First Julian day, it can be present many days, or few days. *The number of days and first Julian day were log_10_ transformed to improve assumptions of normality.*

### Network structural metrics

Before calculating network metrics, we normalized network matrices by dividing each cell value by the number of days the community was observed (zeros remain zeros). We did this because there was unequal sampling effort across studies, even though there was equal effort across networks within studies. Normalizing resulted in non-integer values in some cases, but standardized individuals observed per unit time, therefore the network is weighted by the frequency of interactions for a given monitoring time.

We calculated four species-level network properties: (i) standardized specialization for each species (d*′*) following Blüthgen *et al*. (Blüthgen et al. 2006). We used this measure instead of species degree (number of other species the focal species interacts with), because degree is based on a binary matrix, and so does not utilize information on the frequency of interactions. Specialization (d′) calculates how strongly a species deviates from a random sampling of interaction partners available. We also calculated (ii) interaction strength asymmetry (*ia*), which measures the average mismatch between a focal species’ effect on its interaction partners and the effect of the interaction partners on the focal species (Vazquez et al. 2007). In addition, for each network we identified modules—sets of species that are more connected to each other than to other species in the network (Olesen et al. 2007)— and for each species we calculated (iii) within-module degree (*z*, the standardized number of links per species within a module), and (iv) among-module connectivity (*c*, how well does the species connect different modules). We chose these because we were interested in how phenology affected network metrics, so needed to use metrics that were quantified at the species level (where phenology was varying). In addition, within-module degree (*z*) and among-module connectivity (*c*) characterize the roles that species play in a network, providing a rich way of understanding networks (Olesen et al. 2007, see also Figure 2).

**Figure 1.**
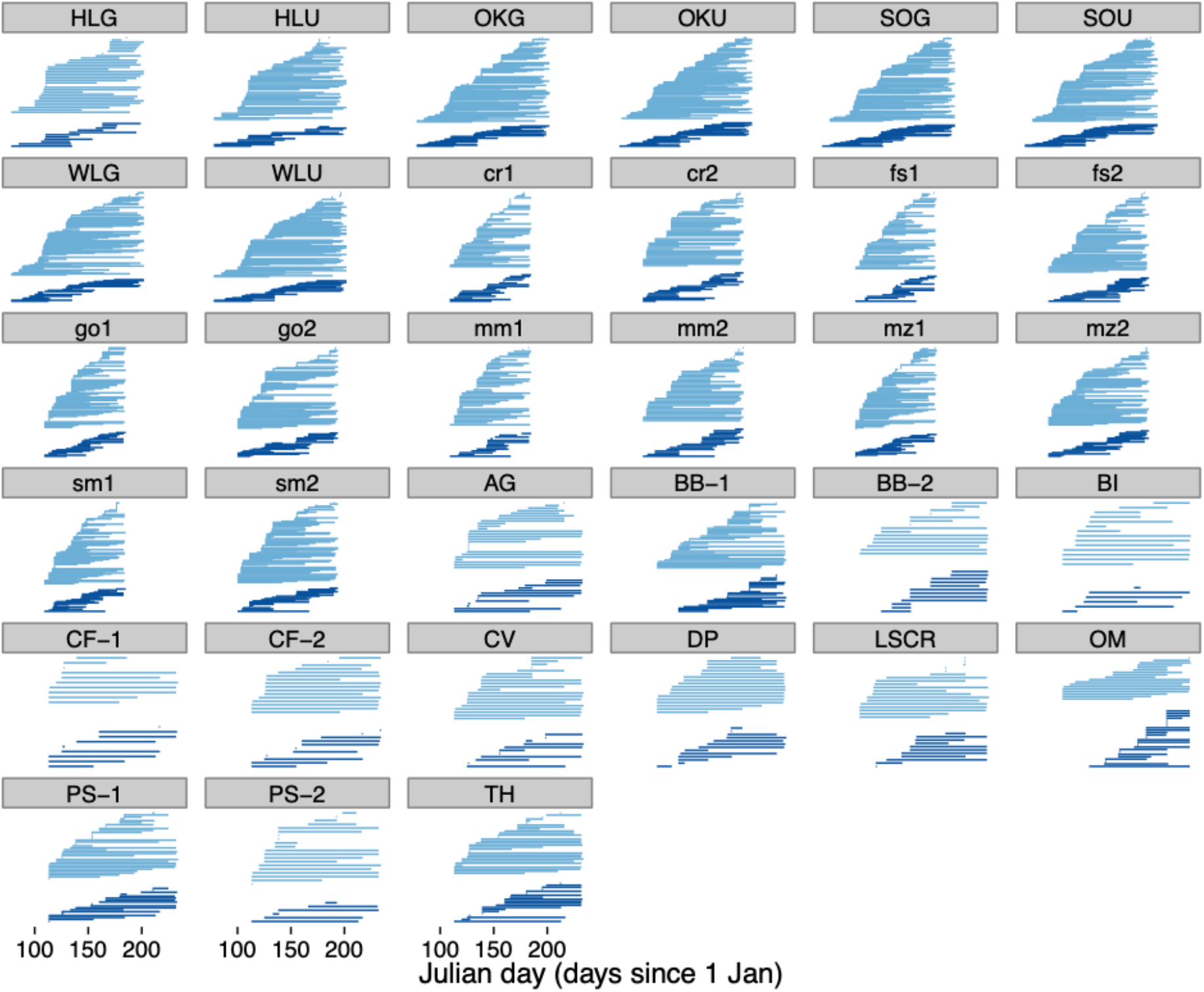
Visualization of the phenology of species in each network, for all 33 networks. The top set of horizontal lines in each panel are pollinators, and the bottom set are plants. Note that each panel follows the same x-axis.

**Figure 2.**
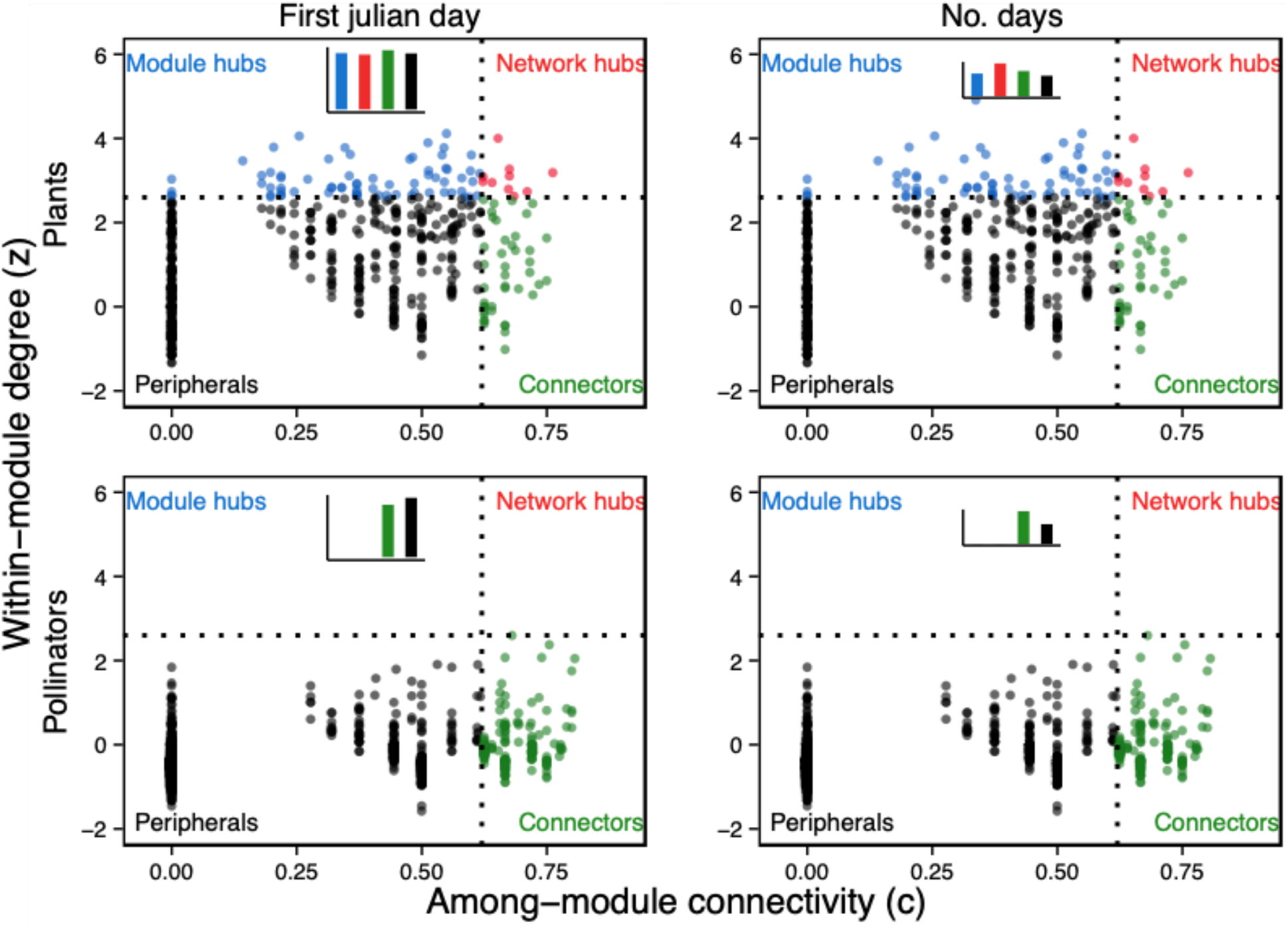
Visualization of species level modularity metrics of within-module degree and among-module connectivity for plants and pollinators. Each panel is split in to four quadrants: Module hubs: species with high z, but low c, or those interacting a lot within their module, but not much among modules; Network hubs: species with both high z and c, or super generalists, acting as both connectors and module hubs; Connectors: species with low z, but high c, or those not interacting a lot within their module, but tending to connect modules; and Peripherals: species with both low z and c, or specialists, i.e,. they have only a few links and mostly within the module. Left-hand panels: bar plots show mean values of first Julian day of appearance in the network for each quadrant. Right-hand panels: bar plots show mean values of total number of days observed in the network for each quandrant. Format following Olesen et al. (2007).

We used the *specieslevel* function in the bipartite R package (Dormann 2011) to calculate d*′*, and *ia*. To calculate the species-level modularity metrics (*z* and *c*), we used a modularity-detecting algorithm, which maximized modularity using simulated annealing (SA) implemented in the command line function *netcarto_cl* in the C program Rgraph (Guimera and Amaral 2005a, 2005b). All other analyses were done with the programming language R (R core team 2018).

### Data analyses

#### How do date of first appearance in a network, and length of activity during the season, affect individual species interaction patterns?

We tested for a relationship between the four species-level network structures (d*′*, *ia*, *c, z*) and phenological variables. For *ia* and *z* (as continuous values from 0 to infinity) we used linear mixed effects models, with two phenology variables (date of first appearance and days observed) as fixed effects and network as a random effect. We also included a random effect for taxonomic group at the family level. This random effect was included as some groups will tend to have a longer duration in the season, for example, bumble bees have multiple generations throughout a season and will therefore be present for a longer period. Including these random effects allows different families to vary either in the intercept or the slopes. We compared five models for each predictor where we varied whether the family random effect was: (i) only for the intercepts (1|family), (ii) a random slope for the number of days but no covariance in between the intercept and slope (0+days|family), (iii) a random slope for the first Julian day with no covariance between the intercept and slope (0+first julian |family), (iv) a random slope for days with covariance between the intercept and slope (1 + days | family) and (v) a random slope for first Julian day with covariance between the intercept and slope (1 | first julian |family). For c and d*′*, variables that are proportions (values between 0 and 1), we used non-linear mixed effects models, with the same formula as above, but specifying a binomial distribution. We selected the best fitting model for the family random effect using AIC (Aho et al. 2014). Models were run separately for plants and pollinators, and for each network metric separately, for a total of eight models. The number of days and first Julian day were log_10_ transformed to improve assumptions of normality and homoscedascity of residuals.

#### How robust are networks to removal of species due to varying phenological “traits”?

We were interested in four scenarios with respect to network robustness: 1) species are removed from a network according to when they are first active – i.e., the species that was first active during the **earliest date** of the season is removed first, and so on; 2) species are removed from a network according to the reverse order of activity – i.e., the species that was first active on the **latest date** is removed first, and so on; 3) species are removed from a network according to total duration of activity during the season – i.e., the species active the **least** number of days is removed first, and so on; and 4) species are removed from a network according to reversed order of total duration of activity during the season – i.e., the species active the **most** number of days is removed first, and so on. These can be referred to as FJ (first julian), FJr (first julian reverse), D (days), and Dr (days reverse), respectively for 1), 2), 3), and 4).

To measure network robustness we used the function *second.extinct* in the *bipartite* R package (Dormann et al. 2009), which works by removing a species, then counting how many species are subsequently lost due to the first removal (i.e., secondary extinctions), and so on for each species removed. This is used to generate a curve of number of species going extinct in one group (plants or pollinators) based on extinctions in the other group. The function *robustness* in the *bipartite* package measures the area under this curve to generate a statistic *R*, with the maximum value *R*=1 corresponding to a very robust network, whereas *R*=0 corresponds to a network that loses species quickly upon initial extinctions.

We ran eight separate simulations, four for each of plants and pollinators, with the four for each being one of the four scenarios outlined above. Each run for each network results in a single robustness (*R*) value. In addition, we calculated *R* for a set of 100 iterations randomly removing species from each network, rather than removing species based on their phenological traits. We compared the “observed” value of *R* for the distribution of each simulation run, and calculated a P-value based on whether the observed value fell outside of the 95% confidence interval of the distribution of *R* from the randomly removed species set.

To compare results between sets of simulation runs (e.g., to compare FJ and FJr within plants), we used two-sample t-tests.

## Results

### Do plants and pollinators differ in their species roles in the network?

A useful way to visualize modularity is plotting among-module connectivity (c) against within-module degree (z) following Olesen et al. (2007). In Figure 2, species fall into roughly defined roles in the network, one of module hubs, network hubs, connectors, or peripherals. To list a few examples of species that fall into one of the four roles, two examples of plant network hubs have generalized flower morphology: *Achillea millefolium* (Asteraceae; *c*=0.65, *z*=4.0), and *Camassia quamash* (Asparagaceae; *c*=0.76, *z*=3.18). Two plant module hubs are the early-flowering *Lomatium macrocarpon* (Apiaceae; *c*=0.60, *z*=2.97), and the late-flowering *Holodiscus discolor* (Rosaceae; *c*=0.14, *z*=3.46). Two examples of pollinator connectors were social bumble bees: *Bombus flavifrons* (Apidae; *c*=0.68, *z*=2.59), *Bombus mixtus* (Apidae; *c*=0.74, *z*=2.1), and two pollinator peripherals were uncommon solitary bees: *Panurginus atriceps* (Andrenidae; *c*=0.28, *z*=1.14), *Osmia tristella* (Megachilidae; *c*=0.28, *z*=1.0). We found that pollinators were neither module nor network hubs.

### How do date of first appearance in a network, and length of activity during the season, affect individual species interaction patterns?

We ranked models based on AICc, and identified top models based on a criteria of ΔAICc < 2.0 from the best model (Steel et al. 2013). On average, species level network metrics were positively related to phenology variables. For both plants and pollinators within-module degree (*z*) and among module connectivity (*c*) were positively related to the number of days in a network (plant and pollinator *P* < 0.001), such that plants and pollinators that were active longer in the season were more connected within the module and among modules (Table 1; Fig 2–3).

**Table 1.**
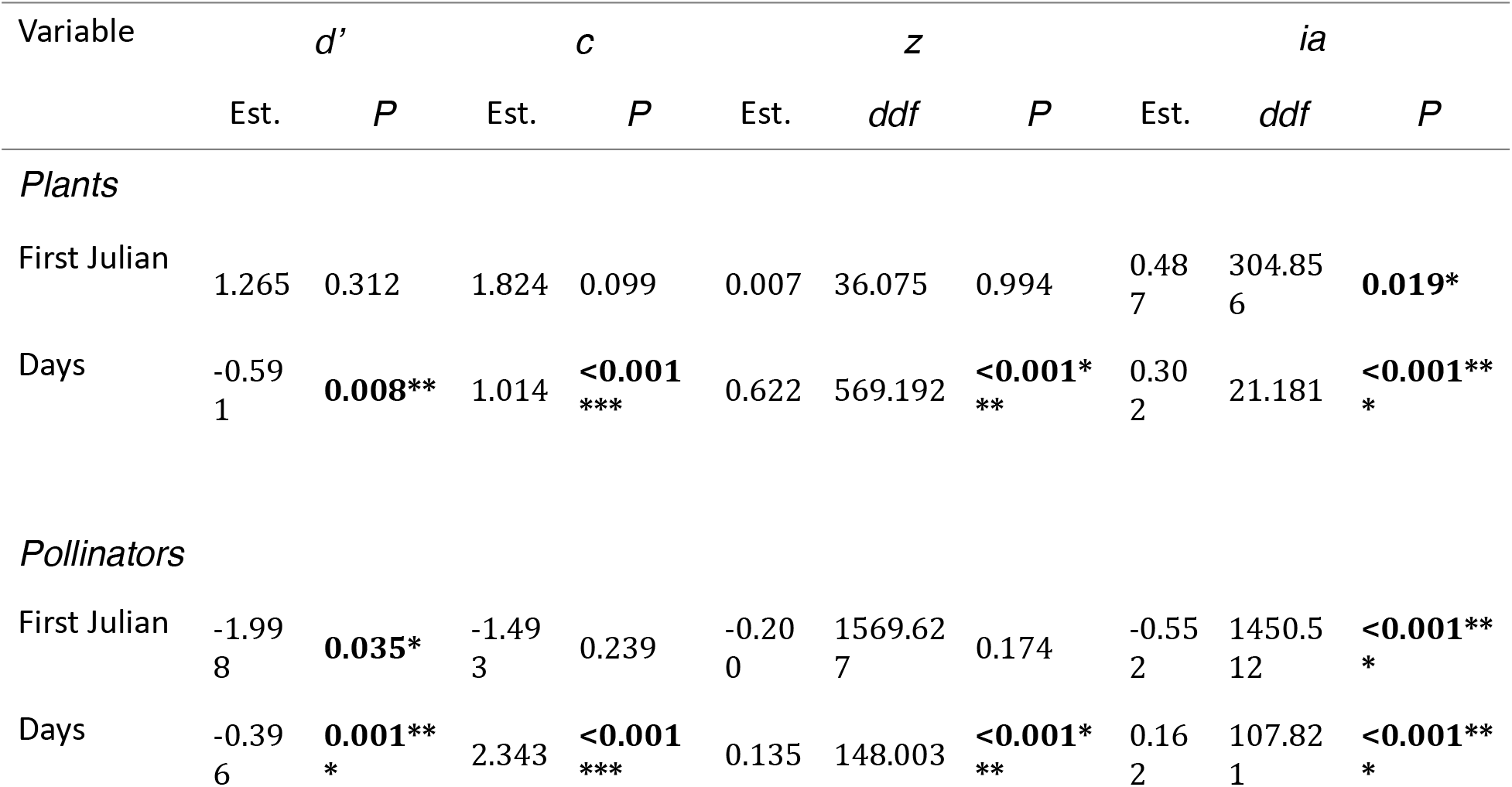
Results of analyses of relationship between two phenology variables (first day of appearance in a network, days observed in a network) and four network structures (specialization (d’), among module connectivity (*c*), within module degree (*z*), and interaction asymmetry (ia)). This table represents eight separate statistical models, one for each of pollinators and plants, and one for each response variable. *, p< 0.05; **, p < 0.01; ***, p < 0.001.

**Figure 3.**
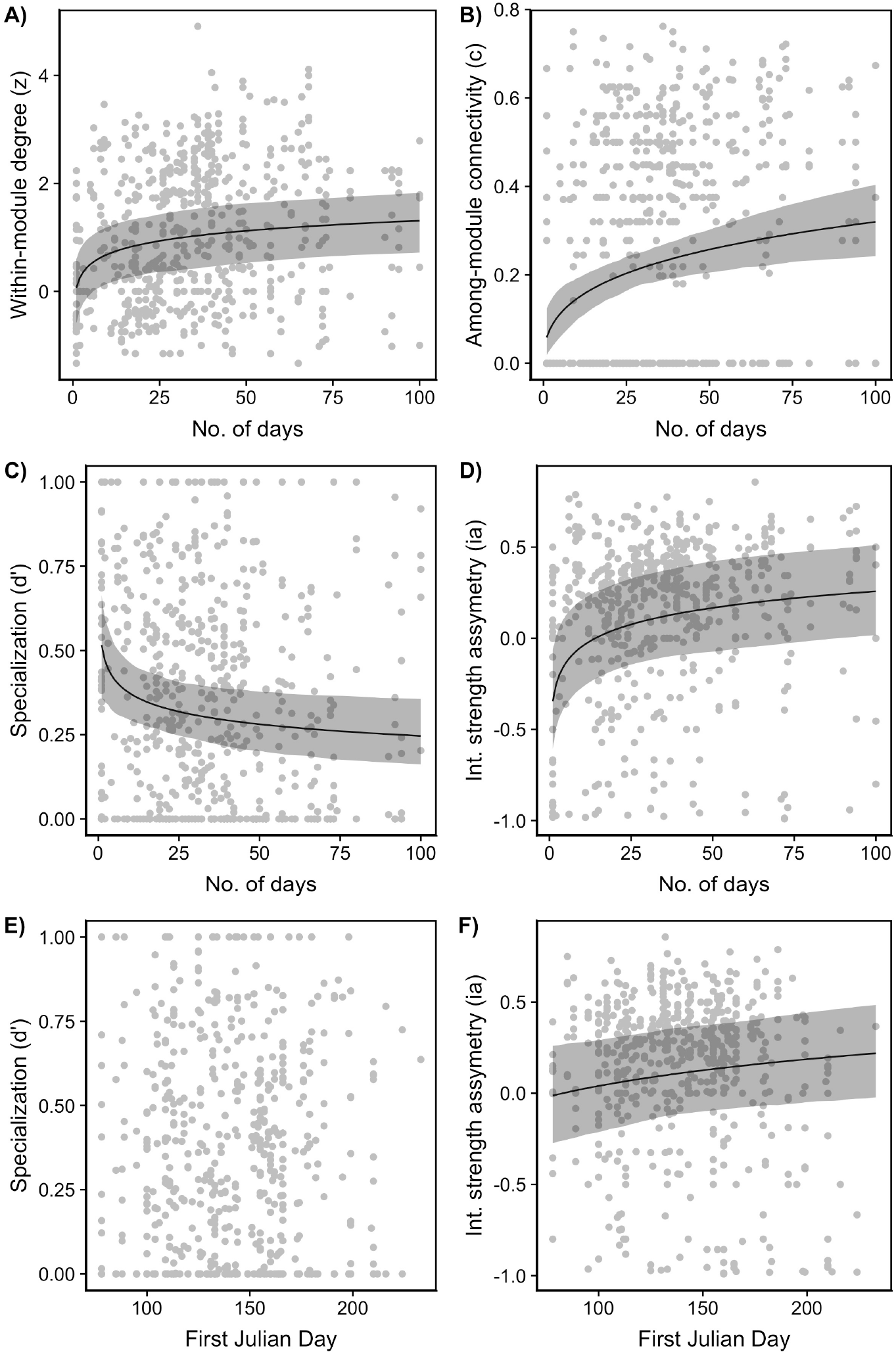
Various network metrics in relation to pollinator and plant phenological traits for plants. A-D) Number of days observed in a network vs. A) within-module degree (z), B) among-module connectivity (c), C) specialization (d’), and D) interaction strength asymmetry (ia). E-F) The date of first appearance in a network vs. E) specialization (d’), and F) interaction strength asymmetry (ia).

The degree of specialization (d’) for plants and pollinators was negatively related with the number of days in a network (*P* = 0.008, *P* = 0.001 respectively), such that plants and pollinators that were active longer in the season were more generalized. For pollinators, the degree of specialization (d’) was negatively related with the day of first appearance (*P* = 0.035), such that pollinators that appeared later in the season were more generalized (Table 1; Figure 2,3,4).

**Figure 4.**
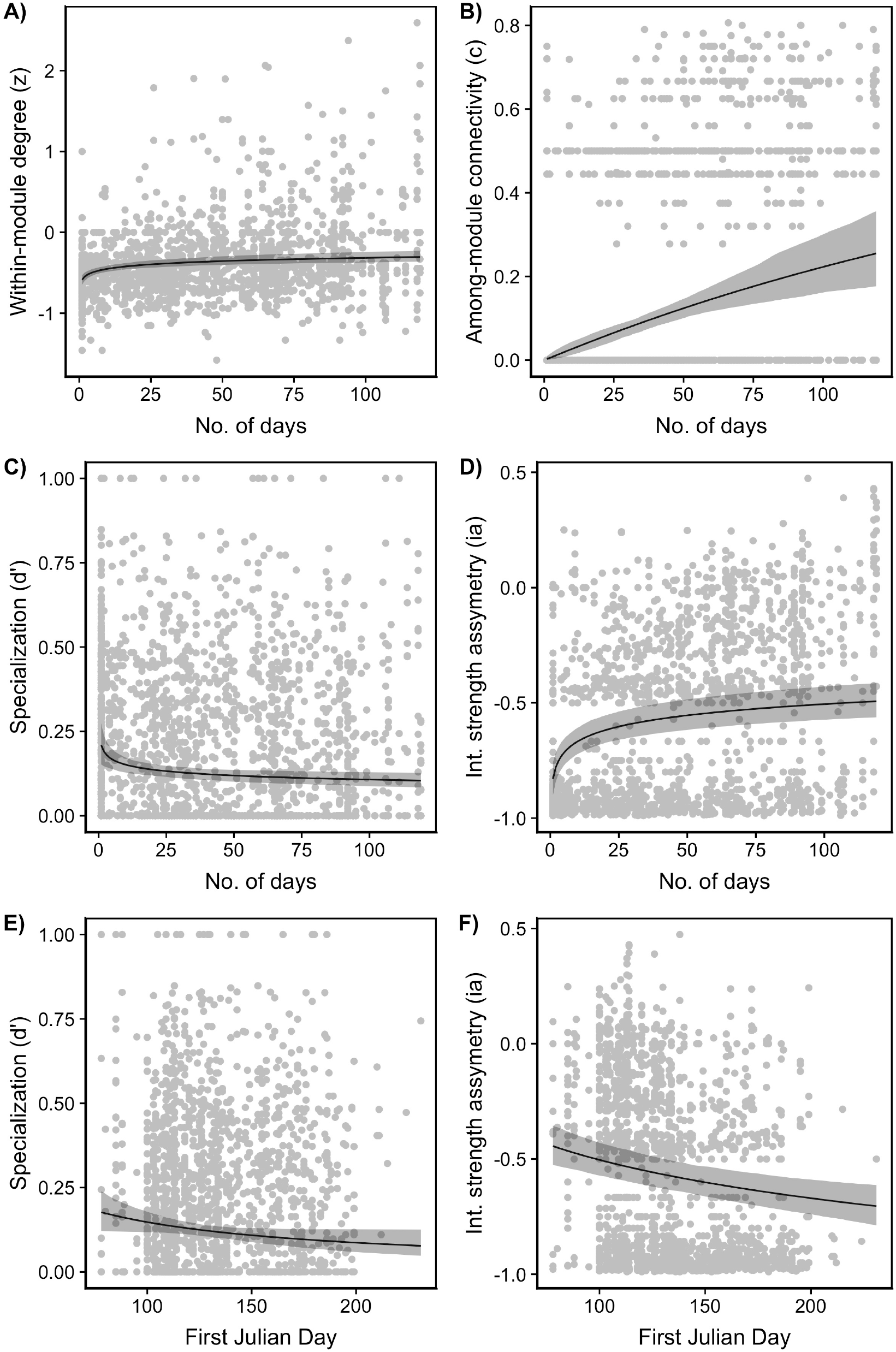
Various network metrics in relation to pollinator and plant phenological traits for pollinators. A-D) Number of days observed in a network vs. A) within-module degree (z), B) among-module connectivity (c), C) specialization (d’), and D) interaction strength asymmetry (ia). E-F) The date of first appearance in a network vs. E) specialization (d’), and F) interaction strength asymmetry (ia).

Last, interaction asymmetry was positively related to the number of days active for both pollinators and plants (plants and pollinators *P* < 0.001), such that plants and pollinators that were active longer in the season affected the species they interacted with more than those species impacted them. The date of first appearance was positively related with interaction asymmetry for plants (*P* = 0.019) and negatively related for pollinators (*P* < 0.001). Plants that appeared later in the season had a stronger effect on pollinators they interact with, while pollinators that appeared early the season had a stronger effect on the plants they interact with. On average pollinators were more strongly affected by the plants they interacted with (Figure 3D and 3F, points below zero) while the plants affected pollinators more strongly (Figure 4D and 4F, points above zero).

### How robust are networks to removal of species due to varying phenological “traits”?

Robustness values (*R*) varied from about 0.40 (less robust) to about 0.92 (more robust; Figure 5). Interestingly, values of *R* were on average higher for pollinator removals than plant removals, suggesting that networks are more robust to removal of pollinators than to removal of plants. For plants, when species were removed first if they appeared early (Figure 5A) in the network 18 of 33 (55%) were significantly different from random species removal. In contrast, when species were removed first if they appeared late in the network, only 9 of 33 (27%) networks were significantly different from random species removal. Overall, network robustness did not differ with removal of plant species appearing early (mean ± 1 s.e.; 0.62 ± 0.01) vs. appearing late (0.59 ± 0.01; Welch’s 2 sample t-test, P = 0.136; n=33 for all comparisons). When species were removed first based on the shortest number of days in a network (Figure 4B) 29 of 33 (88%) were significantly different from random species removal, whereas only 9 of 33 (27%) networks were significantly different from random species removal when species were removed starting with those active the most number of days. Overall, network robustness was greater with removal of species active shortest (0.67 ± 0.01) vs. active longest (0.56 ± 0.01; *P* < 0.001).

**Figure 5.**
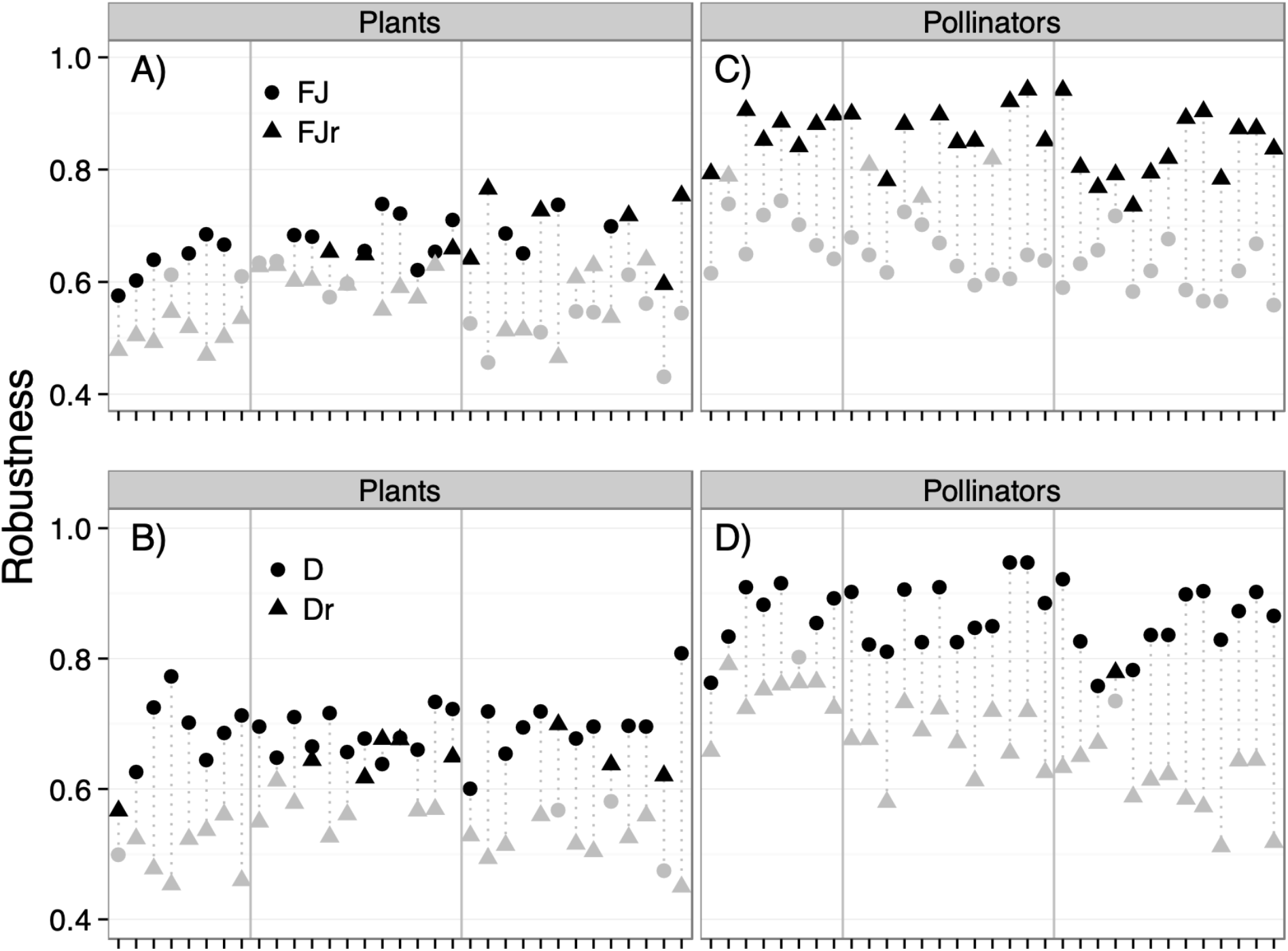
Robustness of 33 plant-pollinator networks in response to removal of species according to either first Julian date of appearance in a network (FJ, circles), last Julian date of appearance (FJr, triangles), least to most days observed in network (D, circles; last day minus first, in number of days), and most to least days observed in network (Dr, triangles; last day minus first, in number of days). Black symbols were significantly more robust than the null model (random species removal), while grey symbols did not differ from the null model. Note: networks in each of the four panels are in the same order as Figure 1; scales in all four panels are the same; drop lines connect points that belong to the same network in each panel.

For pollinators, when species were removed first when they appeared early (Figure 5C) in the network 0 of 33 were significantly different from random species removal, whereas 29 of 33 (88%) networks were significantly different from random species removal with removal starting with species that appeared late in the network. Overall, network robustness differed for those with removal of species appearing early (0.65 ± 0.01) vs. appearing late (0.85 ± 0.01; *P* < 0.001). When species were removed first based on the shortest number of days in a network (Figure 5D) 31 of 33 (94%) were significantly different from random species removal, whereas only 1 of 33 (3%) networks was significantly different from random species removal with removal starting with species that were active the most number of days. Overall, network robustness was greater for those with removal of species active shortest (0.86 ± 0.01) vs. active longest (0.67 ± 0.01; *P* < 0.001).

## Discussion

We asked how species phenological traits (date of appearance in a community, and duration of activity in that community) influenced resulting network structure of 33 plant-pollinator mutualistic networks across three habitat types in British Columbia, Canada. Importantly, we showed in one of the few empirical examinations that phenology, a species attribute that can be sensitive to climate change (Molnár et al. 2012, Bartomeus et al. 2013a), can be an important predictor of structure. This has implications for how mutualistic plant-pollinator networks may respond to climate change via changes in species phenologies.

Species within a module are linked more tightly together than they are to species in other modules. Identifying which species are in these modules can help us understand the mechanisms that structure these networks (Olesen et al 2007). Overall we found that plants were network and module hubs, while pollinators were not. Network hubs —those species that had high within and among network connectivity— and module hubs —species that had high within module but low among module connectivity— tended to be species with radially symmetrical flowers and “easy access” floral rewards. Morphological adaptations that mean flowers can provide food for a diverse array of visiting insect species may increase connectivity within and among modules (e.g. Asteraceae, Asparagaceae, Amaryllidaceae in our study; Wolfe and Krstolic 1999, Sargent 2004).

Pollinators were neither module hubs nor network hubs. This is partly explained by ecological traits of plants and pollinators. For example, pollinator species that were connectors tended to be species that were eusocial (*Bombus*) or likely social (*Lasioglossum (Dialictus) pacatum*) which have long activity times as they have multiple generations per season (Michener 2000). In addition, this result can also be partly explained by a statistical effect, because (as in most network research) the number of pollinators and plants differed greatly. Across all studies, the number of pollinators was three times that of plants. Therefore any given plant is more likely to interact with multiple sets of pollinators than any given pollinator.

### How do date of first appearance in a network, and length of activity during the season, affect individual species interaction patterns?

Our goal was not only to describe species’ network properties (such as modularity), but also, whether phenology was the determinant of those species network properties. Phenology was an important predictor of species-level network structures for both plants and pollinators in this study. This is consistent with findings from numerical simulation studies (Encinas-Viso et al. 2012) that found that phenology could be extremely important in structuring networks. For plants we found that species active longer in the season were more connected both within their module and among modules. This suggests that species are more likely to interact with many different partners the longer they remain active. In addition, plant species that were active longer in the season were also less specialized in their interactions, and had a stronger effect on pollination partners than partners had on them. This suggests that plant species that are active longer during the season are important structurally in the network as they interact with multiple species, exert strong effects on interaction partners and are generalists (Guimarães et al. 2007, González et al. 2010). Some of the species that were active the longest were *Cornus stolonifera (*Family: Cornaceae), *Symphoricarpus albus* (Caprifoliaceae) and *Ranunculus repens* (Ranunculaceae). *Ranunculus* is a weedy forb while *Symphoricarpus* and *Cornus* are a long lasting shrubs. These species additionally have radially symmetrical flowers allowing many types of pollinators to access the flowers (Lovett-Doust et al. 1990, Wolfe and Krstolic 1999, Stewart-Wade et al. 2002, Sargent 2004).

For pollinators, species that were active longer in the season had more connections among and within modules, were less specialized and had a higher interaction strength asymmetry. These results are consistent with the results we found for plants. Some of the species that were active the longest in the season were *Bombus centrals* (Apidae), *Sphaerophoria weemsi* (Syrphidae), and *Lasioglossum pacatum* (Halictidae). These tend to be highly generalized in floral visit patterns (Laverty and Plowright 1988). *Bombus* for example have multiple generations of workers per season (Michener 2000) and although individual workers may specialize on particular floral resources, the species as a whole uses diverse plants over an extended flight period. Hover flies (Syrphidae) also are mutivoltine and use a diverse array of generalized flowers. In contrast, solitary bees in our region are active for only a few weeks, making it unsurprising that they are not within or among module connectors. Similarly, we found that pollinators were less specialized as first Julian day increased. Therefore pollinators that emerged later in the season were more generalist, which in our data would include wasps and some hover flies.

We also found that plant species that emerged later in the season (i.e. their first julian day was higher) had a higher interaction strength asymmetry. Therefore species that emerged later in the season affected their interacting partners more strongly than their partners affected them. Some of the plant species that emerged later in the season were *Galeopsis tetrahit* (Family: Lamiaceae), *Lythrum salicaria* (Lythraceae) and *Polygonum persicaria* (Polygonaceae). Unlike plants, we found that for pollinators the interaction strength asymmetry decreased as species emerged later in the season (i.e. their first julian day was higher). Therefore species that emerged later in the season were more strongly affected by interacting partners than the reverse, and species that emerged earlier in the season affected interacting partners more strongly than partners impacted them. Some of the pollinator species that emerged earlier in the season were solitary bees *Andrena nigrihirta, Andrena merriami, Andrena sladeni, Andrena trizonata and Andrena porterae* (Andrenidae). In two of the ecosystems studied, oak-savannah and shrub-steppe, we observed that at the beginning of the season, many plant species bloom in high density, but temperatures are not yet reliably warm enough for insect activity, so plant reproduction may be pollinator limited (Schemske et al. 1978, Kudo and Ida 2013). By the end of the season the amount of food and nutrients available for the pollinators is lower, but their populations have grown; it may be that late emerging plant species are therefore extremely important resources for pollinators (Mattila and Otis 2007, Garbuzov and Ratnieks 2014). On average, pollinators were more affected by plants (values below 0 on figure 3F and E), while plants affected the pollinators more (values above 0 on figure 4F and 4E). Overall these results suggest that resource limitation may shift along the season from pollinator limitation to flower limitation, a hypothesis that could be tested.

### Robustness

We observed that networks were on average more robust to removal of pollinators than to removal of plants. This makes sense because plants more often link together the plant-pollinator network (hubs organize around plants), whereas pollinators are less often important hubs (see Figure 2). This pattern was also seen in another study – Olesen et al. (2007) found that plant species were more often module hubs and network hubs than pollinators (see Figure 2 in Olesen et al.). The networks we used have 3x as many pollinators than plants. Because of this asymmetry in the number of plants vs. pollinators we would expect that individual plant species would play a more important role than individual pollinators (Vázquez and Aizen 2004).

Removing both plants and pollinators in order from the least to most days observed in the network (D) resulted in more robust networks than removing plants and pollinators in order from the most to least days observed (Dr). This is consistent with our previous results, which found that plants and pollinators that are active the most number of days, were more connected both within and among modules, were less specialized and were affected more strongly by interacting partners. Therefore, removing plants and pollinators that are well connected (i.e. are present the most number of days), results in less robust networks. While this result is expected, the removal of plants and pollinators that are well connected is not significantly different from random removal of species. Therefore, In the face of habitat destruction and climate change, the removal of species due to their duration in the season and the random removal of species could have the same network consequences. Similar simulation studies have found that phenological changes due to climate change can have cascading consequences on the entire network (Revila et al 2015).

Removing plants according to the first Julian date of appearance in a network (FJ) resulted in *more* robust networks than removing plants in order from the last Julian date of appearance (FJr). Plants that appeared later in the season were more important for network robustness than the plants that appeared earlier in the season. On the other hand, removing pollinators according to the first Julian date of appearance in a network (FJ) resulted in *less* robust networks than removing plants in order from the last Julian date of appearance (FJr). Pollinators that appeared early in the season were more important for network robustness than the pollinators that appeared late in the season. These results are consistent with our results for plants and pollinators relating interaction-strength asymmetry to the first julian day of appearance. Again, at the beginning of the season, the system may be more pollinator limited while at the end of the season pollinators may be resource limited (Schemske et al. 1978, Mattila and Otis 2007, Kudo and Ida 2013, Garbuzov and Ratnieks 2014). Our results are consistent with Ramos-Jiliberto et al. (2018), who found that plant persistence was most sensitive to the disappearance of pollinators that start earlier or finish later in the season, and that pollinators were most sensitive to the disappearance of plants that started early and had long seasons.

Plant-pollinator networks are at risk both due to the global decline of pollinators and phenological shifts due to climate change. Pollinator populations are declining globally (Brondizio et al. 2019), specifically bumblebees (*Bombus spp.;* Williams and Osborne 2009, Arbetman et al. 2017). The loss of bumblebees worldwide can have cascading consequences on plant-pollinator networks since bumblebees are active for long periods of time making them connectors in these networks. Losing these connecting species can reduce the robustness of networks as shown in our study. In addition, climate change impacts the phenology of many plant species, affecting in particular spring events such as flowering time (Gordo and Sanz 2010). Earlier flowering time can increase the temporal mismatch between plants and pollinators (Bartomeus et al. 2011, Kudo and Ida 2013). Increasing the temporal mismatch in the spring can further increases the pollen limitation experienced by plants early in the season (Schemske et al. 1978). This mismatch can also reduce the robustness of pollinator networks as shown in our study.

### Conclusion

Our results show that across a large sample of 33 networks, species phenology can be an important predictor of network structure. In particular, the number of days a species is active in a network predicted how connected they are for both plants and pollinators. We also found that plants tended to be more important in the network at the end of the season, while pollinators were important at the beginning of the season. Future work should build on the work presented here by exploring how experimental or natural changes in phenological variables, like time of first appearance or duration of activity in a community, influence network structure.

## Acknowledgements

We thank the Elle lab group for comments that improved earlier versions of this manuscript. S. Elwell, G. Gielens, and H. Gehrels kindly shared data. We acknowledge funding from NSERC-CANPOLIN, the Canadian Pollination Initiative, and an NSERC Discovery Grant to EE.

