## Supplementary material for "Phenology and Robustness in plant-pollinator networks": Table A1

Appendix A. Details for each of the study sites.

Table A1. Details of each of the study sites in British Columbia, Canada. Abbrevations: oak sav. = oak savannah; antel. steppe = antelope shrub steppe; sage. steppe = sagebrush steppe.

| Site name (abbrev.) | Year(s) Coll. | Ecosystem | Latitude | Longitude |
|---|---|---|---|---|
| go1, go2 | 09/10 | oak sav. | 48.808 | -123.631 |
| mz1, mz2 | 09/10 | oak sav. | 48.791 | -123.638 |
| fs1, fs2 | 09/10 | oak sav. | 48.819 | -124.134 |
| mm1, mm2 | 09/10 | oak sav. | 48.818 | -124.117 |
| sm1, sm2 | 09/10 | oak sav. | 48.782 | -123.885 |
| cr1, cr2 | 09/10 | oak sav. | 48.777 | -123.941 |
| HLU | 10 | antel. steppe | 49.087 | -119.518 |
| HLG | 10 | antel. steppe | 49.091 | -119.527 |
| OKU | 10 | antel. steppe | 49.262 | -119.509 |
| OKG | 10 | antel. steppe | 49.184 | -119.586 |
| WLU | 10 | sage. steppe | 49.303 | -119.630 |
| WLG | 10 | sage. steppe | 49.312 | -119.680 |
| SOU | 10 | sage. steppe | 49.015 | -119.588 |
| SOG | 10 | sage. steppe | 49.011 | -119.621 |
| AG | 13 | urban forest | 49.010 | -122.451 |
| BB-2 | 13 | urban forest | 49.017 | -123.052 |
| BI | 13 | urban forest | 49.175 | -122.580 |
| CV | 13 | urban forest | 49.018 | -122.664 |
| BB-1 | 13 | urban forest | 49.016 | -123.041 |
| CF-1 | 13 | urban forest | 49.241 | -122.809 |
| CF-2 | 13 | urban forest | 49.240 | -122.796 |
| DP | 13 | urban forest | 49.079 | -122.938 |
| LSCR | 13 | urban forest | 49.395 | -122.992 |
| OM | 13 | urban forest | 49.238 | -123.126 |
| PS-2 | 13 | urban forest | 49.254 | -123.196 |
| PS-1 | 13 | urban forest | 49.271 | -123.259 |

| Site name (abbrev.) | Year(s) Coll. | Ecosystem | Latitude | Longitude |
|---|---|---|---|---|
| TH | 13 | urban forest | 49.184 | -122.750 |
