## Supplementary material for "Phenology and Robustness in plant-pollinator networks": Table A2

### Table S1

Model structure was: response ~ log10(first_julian) + log10(days) + (1 | pi/.id) + Family random effect.

***Plants***

| Model | Family random effect | z | | c | | d’ | | ia | |
|---|---|---|---|---|---|---|---|---|---|
| | | df | AIC | df | AIC | df | AIC | df | AIC |
| 1 | (1 \| FamilyName) | 7 | 1848.628 | 6 | 606.699 | 6 | 750.7722 | 7 | 517.8107 |
| 2 | (0 + log10(days) \| FamilyName) | 7 | 1848.566 | **6** | **606.699** | **6** | **748.7950** | 7 | 530.1231 |
| 3 | (0 + log10(first_julian) \| FamilyName) | 7 | 1849.441 | 6 | 606.699 | 6 | 751.2407 | 7 | 517.7547 |
| 4 | (1 + log10(days) \| FamilyName) | 9 | 1847.378 | 8 | 610.699 | 8 | 752.1878 | **9** | **505.4113** |
| 5 | (1 + log10(first_julian) \| FamilyName) | **9** | **1845.088** | 8 | 610.699 | 8 | 755.2375 | 9 | 520.9011 |

***Pollinators***

| Model | Family random effect | z | | c | | d’ | | ia | |
|---|---|---|---|---|---|---|---|---|---|
| | | df | AIC | df | AIC | df | AIC | df | AIC |
| 1 | (1 \| FamilyName) | 7 | 2027.596 | 6 | 1099.950 | 6 | 1332.472 | 7 | 1105.336 |
| 2 | (0 + log10(days) \| FamilyName) | **7** | **1996.510** | 6 | 1099.950 | **6** | **1332.472** | **7** | **1072.677** |
| 3 | (0 + log10(first_julian) \| FamilyName) | 7 | 2030.213 | 6 | 1099.950 | 6 | 1332.472 | 7 | 1108.556 |
| 4 | (1 + log10(days) \| FamilyName) | 9 | 1998.208 | **8** | **1089.255** | 8 | 1336.472 | 9 | 1076.677 |
| 5 | (1 + log10(first_julian) \| FamilyName) | 9 | 2017.617 | 8 | 1103.950 | 8 | 1336.472 | 9 | 1090.839 |
